# A minimally sufficient model for rib proximal-distal patterning based on genetic analysis and agent-based simulations

**DOI:** 10.1101/179077

**Authors:** J. L. Fogel, D. L. Lakeland, I. K. Mah, F. V. Mariani

## Abstract

For decades, the mechanism of skeletal patterning along a proximal-distal axis has been an area of intense inquiry. Here we examine the development of the ribs, simple structures that in most terrestrial vertebrates consist of two skeletal elements— a proximal bone and a distal cartilage portion. While the ribs have been shown to arise from the somites, little is known about how the two segments are specified. During our examination of genetically modified mice, we discovered a series of progressively worsening phenotypes that could not be easily explained. Here, we combine genetic analysis of rib development with agent-based simulations to conclude that proximal-distal patterning and outgrowth could occur based on simple rules. In our model, specification occurs during somite stages due to varying Hedgehog protein levels, while later expansion refines the pattern. This framework is broadly applicable for understanding the mechanisms of skeletal patterning along a proximal-distal axis.

## INTRODUCTION

During evolution, a number of changes in vertebrate body plan allowed terrestrial tetrapod species to thrive on land and take advantage of new habitats. For example, in contrast to the open rib cages of aquatic species, the thoracic cage became an enclosed chamber that could support the weight of the body and facilitate lung ventilation (Janis and Keller, 2001). Current tetrapod species typically have ribs that are subdivided into two segments, a proximal endochondral bony segment connected to the vertebrae, and a distal permanent cartilage segment that articulates with the sternum (Fig. 1A, B). The rib bones support the body wall and protect the internal organs; the costal cartilage maintains thoracic elasticity, allowing respiration while still enclosing the thoracic cage. Although clues from the fossil record are beginning to reveal when the enclosed rib cage arose during evolution (Daeschler et al., 2006; Pierce et al., 2013), little is known about what changes occurred during embryogenesis to extend the ribs around the body and to connect the ribs to the sternum via a costal cartilage element (Brainerd, 2015). Here using genetic and computational approaches, we generate a plausible model for how two rib segments form during development.

**Figure 1:**
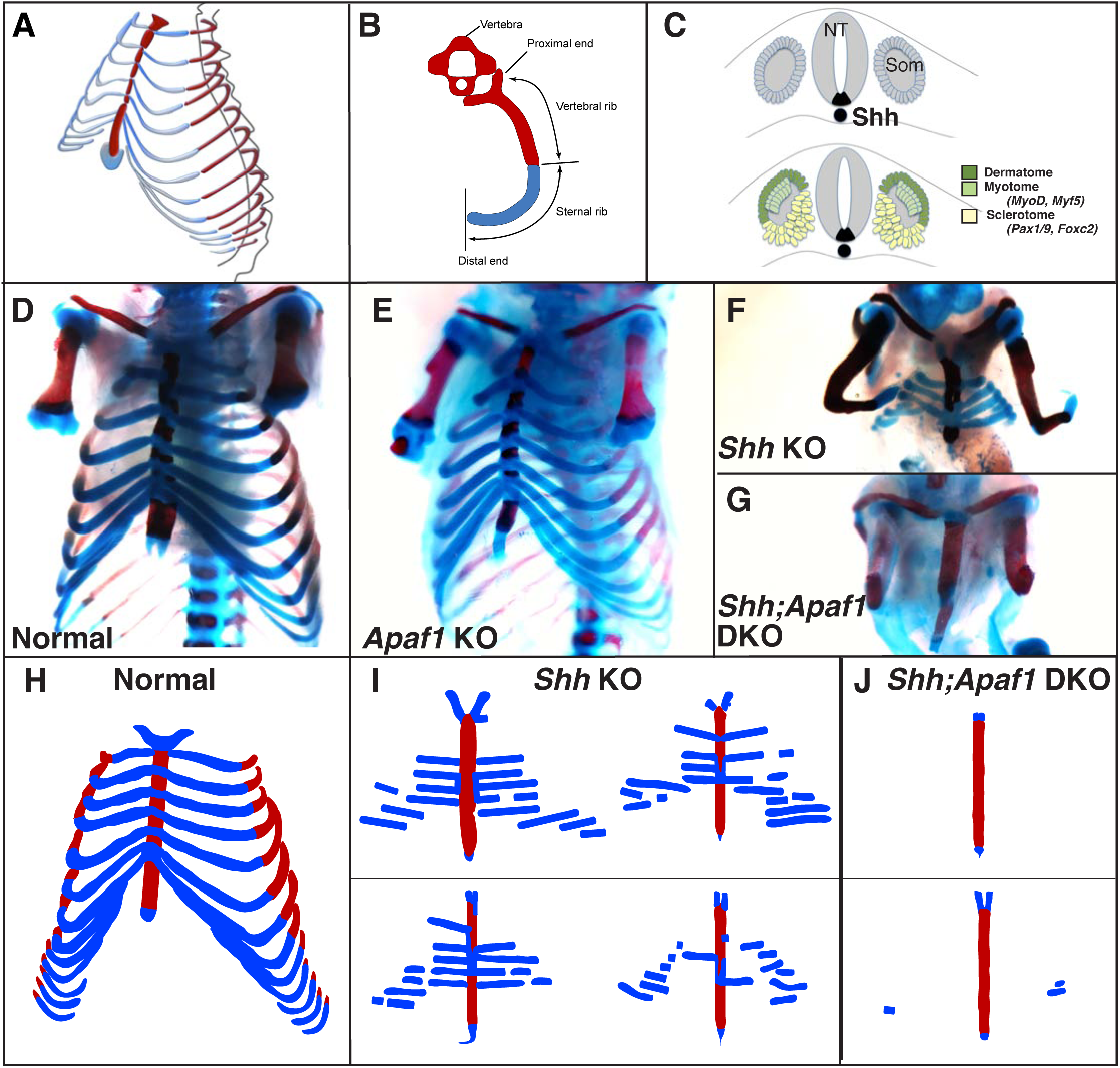
Rib skeletal development is compromised in *Shh* null animals. A. Frontal 3/4 view of the thoracic cage depicting the orientation of the proximal and distal ribs. Mice have 13 pairs of ribs. B. Schematic of a vertebra and rib, transverse view. Red represents bone including the proximal vertebral rib and blue represents the cartilaginous distal/sternal rib. C. The somite (Som), neural tube (NT), and notochord diagramed in cross-section. The dermatome and myotome (dark and light green) gives rise to the dermis and muscles while the sclerotome (yellow) gives rise to the vertebrae and ribs. Markers for these compartments are indicated. The location of *Shh*-expressing cells is indicated in black. **D-G.** Alcian blue and alizarin red skeletal staining of the rib cage: **D.** Normal mouse skeletal development at E18.5 (lower arms removed). **E.** Without *Apaf1*, the axial skeleton develops normally (n=12, lower arms removed). **F.** Without *Shh*, embryos develop without vertebrae and the proximal portion of the ribs. Distal ribs form but are not patterned correctly (n=17). **G.** Double knock out (DKO) of both *Shh* and *Apaf1* results in a more severe phenotype. DKO neonates develop without vertebrae, proximal *and* distal ribs (n=7). The sternum is still present and ossifies on schedule. **H-J.** Schematics representing skeletal preparations of normal (**H**) and null neonates. **I.** The loss of the proximal ribs is consistent amongst all *Shh* KO neonates however the disrupted pattern of the distal ribs vary. **J.** Occasionally *Shh;Apaf1* DKO neonates had cartilage nodules laterally (presumably at the chondro-costal joint, n=1/7).

Lineage tracing studies indicate that the sternum and ribs have different developmental origins. The sternum, like the appendicular skeleton, arises from the lateral plate mesoderm (Cohn et al., 1997; Bickley and Logan, 2014), while the ribs and vertebrae arise from the somites (reviewed in (Brent and Tabin, 2002)). Studies using chicken-quail chimera grafts have shown that the thoracic somites contribute to all portions of the ribs (Huang et al., 1994), with a the medial somite contributing to the proximal ribs while lateral somite contributes to the distal ribs (Olivera-Martinez et al., 2000). These results suggest that the proximal and distal progenitor populations of the rib are separated early on rather than being mixed. As the whole somite matures, it separates into distinct dorsal (dermomyotome and myotome) and ventral (sclerotome) compartments (Fig. 1C). Initially there was some debate on the precise embryological origin of the ribs within the somite (Kato and Aoyama, 1998; Huang et al., 2000). However, using retroviral lineage labeling which avoids the challenges of transplantation experiments, both the proximal and distal segments of the rib were shown to arise from the sclerotome compartment (Evans, 2003). It has been still unclear though, how the sclerotome becomes patterned along the proximal-distal axis.

Through studies particularly of Drosophila wing/leg disc and of vertebrate limb development over the past decades, several patterning models have been conceived to explain how proximal-distal, dorsal-ventral, and anterior-posterior pattern arises (Briscoe and Small, 2015). For example, compartments could become specified based on: 1. the presence of cellular determinants, 2. the concentration of a morphogen, 3. the duration of exposure to a signaling molecule, and/or 4. the action of local relay or mutual inhibition signaling. Specification could gradually emerge over the course of organogenesis or via a biphasic process with specification occurring early in a small population of cells followed later by expansion into compartments (recently reviewed in (Zhu and Mackem, 2017)). In this study, we first use genetically modified mice in which the Hedgehog (Hh) and apoptosis pathway is disrupted to provide clues for how two rib segments are patterned and grow. Our experiments produced unexpected results, which led us to seek an explanation using Agent-Based Modeling, a simulation method based on a cell’s ability to make decisions in response to stimuli. We designed a set of simple rules that could produce a wide variety of potential phenotypes. By adjusting the parameters, we were then able to reproduce the qualitative characteristics seen in our genetically modified mice using several different parameter combinations. In order to eliminate some possibilities and improve our model, we were then motivated to collect further biological measurements. Using a refined model, we were then able to conclude that complex patterning and growth can emerge through a set of simple rules and biologically supported parameters. Furthermore, our model does not require individual cells to have necessarily received any positional information. Finally, we find that our model is essentially biphasic, with early events that define the size and fate of the progenitor populations and later events that expand the already specified population.

Studying rib formation during embryogenesis, and in particular, determining how the segments of the rib cage become distinct provides a relatively simple test-case for questions regarding skeletal patterning, while also giving clues as to the evolutionary origin of the enclosed rib cage. In addition, our studies may aid in understanding the etiology of congenital abnormalities of the rib cage (Blanco et al., 2011). The use of agent-based modeling provides insight into how simple decision making at the cellular level could lead to the emergence of multi-component structures during skeletal development.

## RESULTS

### Impact of blocking cell death in *Shh* null embryos

Previous studies have demonstrated the importance of Hedgehog (Hh) signaling for sclerotome induction and specification. Overexpression of Sonic hedgehog (Shh) can produce ectopic sclerotome at the expense of dermomyotome. Furthermore, in the absence of *Shh* there is a reduction and later loss of the sclerotome marker *Pax1* and a complete loss of vertebrae and the proximal rib segments (Fan and Tessier-Lavigne, 1994; Chiang et al., 1996; Marcelle et al., 1999). The distal cartilaginous portions of the ribs are still present although abnormal. Hh signaling is also well-known to be important for promoting cell proliferation, growth, and survival (Charrier et al., 2001; Thibert et al., 2003) and in the absence of Shh or Shh-producing cells in the floor plate and notochord the number of somite cells going through apoptosis is greatly increased (Teillet et al., 1998). Thus, in addition to a role for Hh signaing in somite/sclerotome induction, Hh signaling also protects somite cells from undergoing programmed cell death. To distinguish between these two roles, we decided to block cell death in *Shh* null animals. In previous studies where loss of *Shh* results in high cell death in the developing heart and face, removing the function of Apoptotic protease activating factor 1 (Apaf1), a central player in the mitochondrial pathway of programmed cell death, could block cell death and rescue the phenotype (Aiyer et al., 2005; Long et al., 2013). Perhaps similarly, removing *Apaf1* on a *Shh* null background would inhibit somite cell death and rescue the thoracic phenotype. We therefore generated genetically modified mouse lines that lacked *Shh*, *Apaf1*, or both.

Apaf1 is required for the Cytochrome c and ATP-dependent activation of Caspase9 which leads to the subsequent activation of Caspase3, followed by the initiation of nuclear breakdown and proteolysis (Zou et al., 1997). Embryos carrying null alleles for *Apaf1* (*Apaf1* KO) have vastly decreased cell death in the nervous system and exhibit disruptions of the head and face skeleton likely due to a grossly overgrown CNS. In addition, interdigital death in the autopod is delayed (Cecconi et al., 1998; Yoshida et al., 1998). Although, the embryos develop to term, they typically die at birth and have not been examined in detail during skeletal development. Therefore, we first investigated *Apaf1* KO skeletal preparations from E14.5-E18.5. We found that patterning of the axial and appendicular portions of the skeleton is grossly normal and that cartilage and bone development proceeds on schedule, indicating that *Apaf1* is not required for normal skeletal development (Fig. 1D, E).

As previously observed, *Shh* KO animals exhibit a complete loss of the vertebral column and pronounced rib cage defects (Fig. 1F) (Fan and Tessier-Lavigne, 1994; Chiang et al., 1996). Loss of the proximal ribs was consistently observed amongst all *Shh* KO embryos stained for bone and cartilage (E15-18). In contrast, the pattern of the remaining rib segments varied (Fig. 1I). These cartilage portions were distally located and never mineralized suggesting that they represented the distal costal cartilages. These segments were not entirely normal as they were discontinuous, reduced in number (∼7-8 instead of 13), not properly articulated with the sternum, and positioned at abnormal angles. The clavicle and sternum were present although the sternum was sometimes not completely fused and had missing or disorganized segments; however it did undergo ossification on schedule. In a few cases, small condensations could be observed laterally, possibly near the chondro-costal joint (Fig. 1F).

We then created animals double null for *Shh* and *Apaf1* (*Shh;Apaf1* DKOs) and analyzed the skeletal pattern. To our surprise, instead of observing a rescue of the *Shh* KO phenotype, *Shh*;*Apaf1* DKO animals exhibited an even more severe skeletal phenotype. *Shh;Apaf1* DKO embryos did display features of the *Shh* single KO (loss of the vertebrae and proximal ribs). However, in addition to these defects, the distal portion of the ribs was now missing as evidenced by the lack of alcian blue staining in the body wall (Fig. 1G). Rarely a few small pieces of cartilage were present at the lateral margin, near the expected location of the chondro-costal joint (Fig. 1J).

### Similar synergistic loss of the axial skeleton with removal of *Caspase3*

Apaf1 has been shown to have other roles in addition to regulating the programmed cell death pathway (Zermati et al., 2007; Ferraro et al., 2011). Thus to determine if the observed effects were specific to Apaf1 or more generally due to altering the cell death pathway, we carried out mouse crosses utilizing a *Caspase3* null allele (*Casp3* KO). Caspase3 is an executioner caspase, that cleaves key structural proteins leading to DNA fragmentation and membrane blebbing (Fuchs and Steller, 2011). As with loss of *Apaf1*, the axial skeleton of *Casp3* KO animals was grossly normal (Fig. 2A,B). *Shh*;*Casp3* DKO embryos exhibited a complete loss of all vertebrae, vertebral ribs and sternal ribs as was observed in the *Shh*;*Apaf1* DKO embryos (compare Fig. 1G and Fig. 2D). These results suggest that the observed phenotypes when *Apaf1* is removed are not due to a specific non-canonical function but rather due to a general abrogation of the programmed cell death pathway.

**Figure 2:**
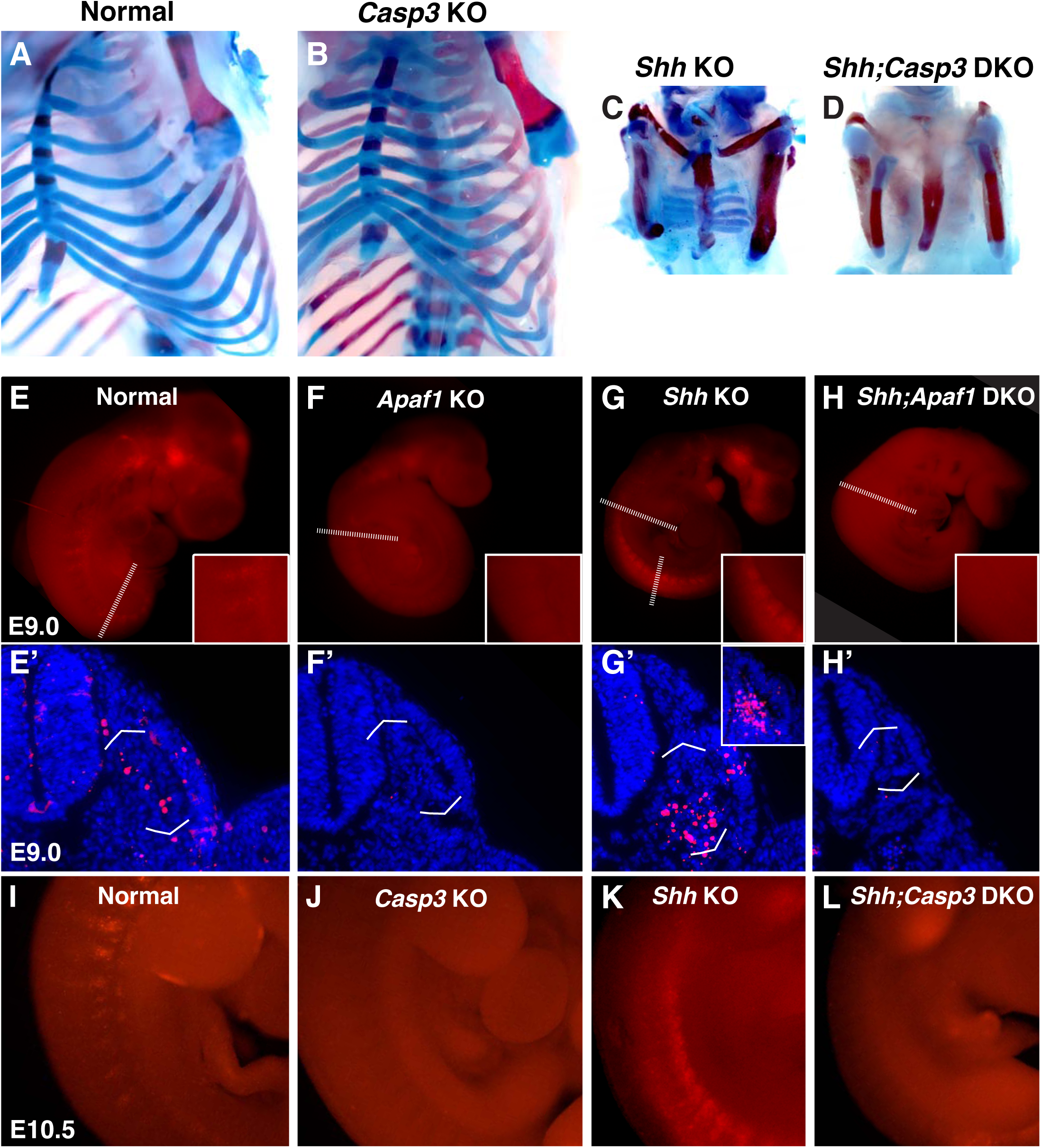
Programmed cell death is decreased in the absence of *Casp3* and *Apaf1*. **A-D.** Alcian blue and alizarin red staining, E18.5 neonates. **A, B.** Normal and *Casp3* KO embryos. **C.** *Shh* KO embryos develop without vertebrae and proximal ribs and as before, distal ribs are present but mispatterned. **D.** *Shh;Casp3* DKO embryos exhibit a similar phenotype to *Shh;Apaf1* DKO embryos— a complete loss of all vertebrae, proximal, and distal ribs. **E-L.** Embryos stained with LysoTracker (red) are shown in whole-mount and in section (blue, DAPI). **E, E’.** Normal embryos have LysoTracker staining in the roof plate, limbs, and in the developing somites. **F, F’.** Throughout *Apaf1* KO embryos, including the somites, LysoTracker staining is dramatically reduced. **G, G’.** *Shh* null embryos have increased LysoTracker staining in the somites. At the thoracic level, staining is found in sclerotomal cells emerging from the ventral medial somite. At a more caudal level (inset) abundant staining is present in the sclerotome epithelium. **H, H’**. *Shh;Apaf1* DKO embryos have little to no LysoTracker staining throughout. **I-L.** E10.5 embryos from the *Shh-Caspase3* cross had similar LysoTracker staining patterns. **I.** Normal embryos. **J.** *Casp3* KO embryos displayed a dramatic reduction in LysoTracker staining. K. *Shh* KO embryos have abundant staining in the somites. **L.** Reduced LysoTracker staining. (similar to *Shh;Apaf1* DKO) was also observed in *Shh;Casp3* KO null embryos. *Phenotypes were consistently observed in at least three animals of each genotype.*

### *Shh;Apaf1* DKO and *Apaf1* KO embryos exhibit a reduction in cell death

To determine if the loss of programmed cell death genes was indeed preventing cells from dying, TUNEL and/or LysoTracker assays were performed (Fogel et al., 2012). In control embryos, evidence of cell death can be seen as distinct periodic stripes of staining in the somites (Fig. 2E, E’, I) a pattern that had been observed previously concentrated in the sclerotome portion beginning as early as E9.0, extending to E10.5 and waning by E11.5 (Sanders, 1997; Teillet et al., 1998). Loss of *Apaf1* however, results in a dramatic reduction in cell death throughout the embryo, and notably absence of cell death in the somites (Fig. 2F, F’). In contrast, *Shh* KO embryos exhibit high cell death in the somites — with high LysoTracker positive staining in the ventral sclerotome domain (Fig. 2G, G’ and inset, marked with brackets). Interestingly, the loss of *Apaf1* in *Shh* null embryos *(Shh;Apaf1* DKOs) results in a vast reduction in LysoTracker positive cells (Fig. 2H, H’). Like *Apaf1* KO embryos, *Casp3* KO embryos display a dramatic reduction in cell death compared to the normal pattern (Fig. 2I, J). Similarly, *Shh*;*Casp3* DKO embryos (Fig. 2L), also exhibit a dramatic reduction in cell death compared to the *Shh* KO pattern. Thus, loss of either *Apaf1* or *Casp3* results in an inhibition of programmed cell death even in *Shh* null animals. However, it seems counterintuitive that a reduction in cell death could lead to a more severe skeletal phenotype.

### *Shh* expressed in the floor plate and notochord is necessary for rib development

Shh has been shown to play an important role in the patterning of a number of different tissues and organs during embryonic development. Previous work has demonstrated that *Shh* expression in the notochord and floor plate is essential for ventral neural tube and somite specification (Varjosalo and Taipale, 2008). However, *Shh* is also expressed in other developing tissues that could affect somite patterning and rib formation. For example, RNA *in situ* hybridization reveals that *Shh* is expressed not only in the notochord and floor plate but also the dorsal root ganglia, ventral neural tube, developing lungs, as well as the developing ribs themselves (Fig. 3A, B). To determine if *Shh* from the notochord and floor plate specifically is required to obtain the observed phenotypes, *Shh* hypomorph embryos were created utilizing a tamoxifen inducible *Foxa2-*CRE conditional knock-out approach (Park et al., 2008). Using this system the temporal influence of Shh on rib development could also be analyzed. Administration of tamoxifen at E8.0 did not alter *Ptch1* expression, a readout of *Shh* signaling, in E9 somites (data not shown) indicating that the activity of Shh prior to cre-mediated deletion (likely sometime between E8.0-E10) was sufficient for *Ptch1* expression. As a result, these embryos had no skeletal defects in the thoracic cage (data not shown, later injections also failed to generate skeletal defects). However, administration of tamoxifen at E7.0 caused a discontinuity in *Shh* and *Ptch1* expression along the ventral neural tube and reduced *Ptch1* in the somites by E8.0 (data not shown). Subsequently, embryos developed with a range of *Shh* KO hypomorphic phenotypes. Importantly, the most severe phenotypes were very similar to *Shh* KO animals and lacked the vertebrae and proximal ribs (Fig. 3C). The distal-most sternal ribs were present but mis-patterned, although less severely than the *Shh* KO animals. Furthermore, the additional loss of *Casp3* again resulted in failed distal rib development (Fig. 3D). Among the hypomorphic *Shh* KO animals, less severely affected animals lacked vertebrae and the proximal half of the vertebral rib, while the least affected only had abnormal vertebrae and were missing just the most proximal ends of the proximal rib (Fig. 3F-H). Thus, these experiments demonstrate that Shh signaling from the notochord and floor plate is essential for normal thoracic skeletal development and is required prior to rib outgrowth and potentially even earlier; furthermore, the additional loss of a cell death gene exacerbates the phenotype. In addition, a comparison of the milder phenotypes vs. the more severe phenotypes suggests that high levels of Hedgehog signaling are required for normal proximal development and lower levels for distal development.

**Figure 3:**
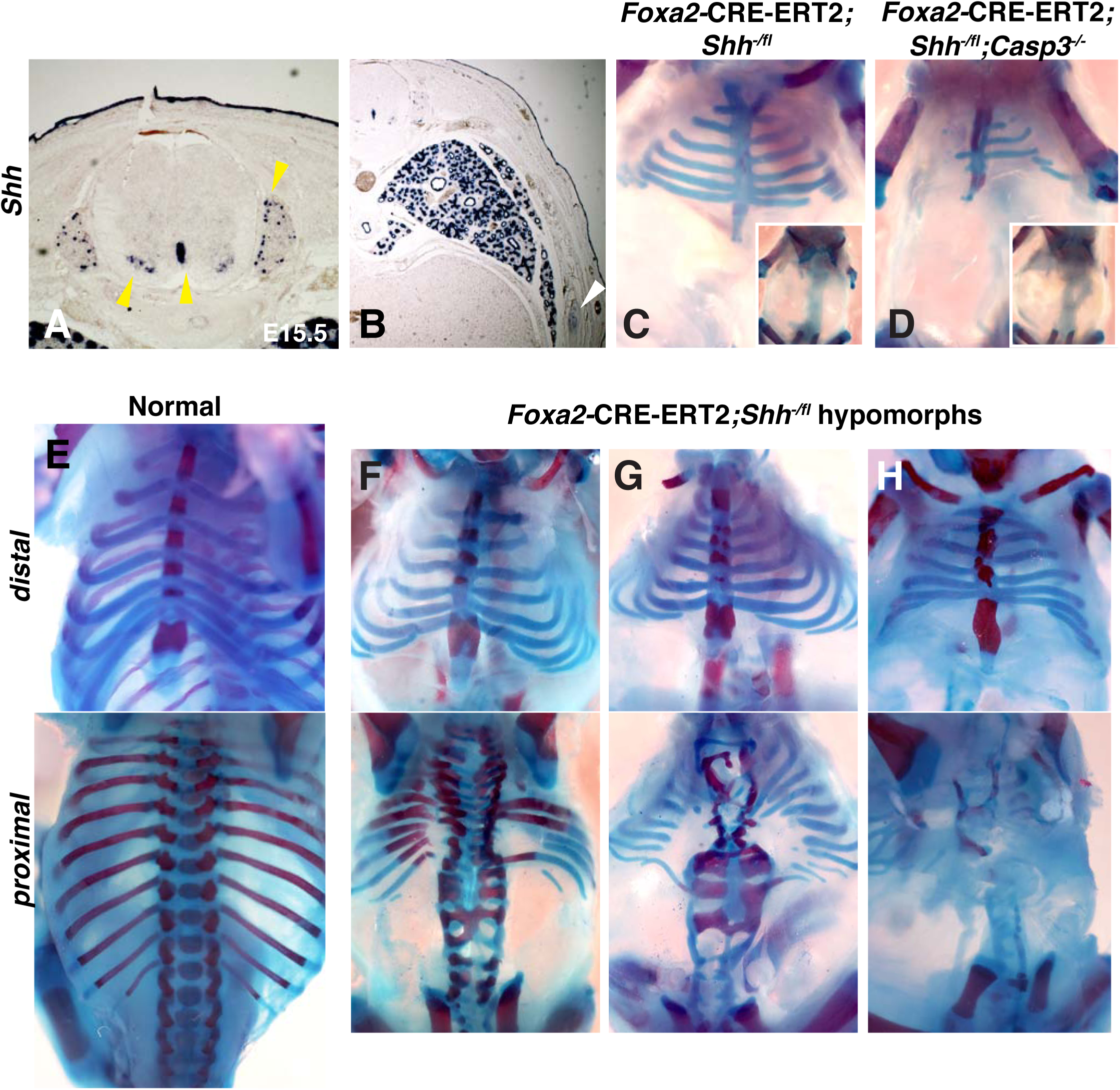
Conditional removal of *Shh* shows an early role for Hh signaling in rib development. **A, B.** *Shh* is expressed not only in the floor plate but also the dorsal root ganglia, ventral neural tube, developing lungs (yellow arrow-heads), as well as the developing ribs themselves (white arrowhead). **C-H**. Alcian blue and alizarin red staining of neonates in which an inducible *Foxa2-CRE-ERT2* was used to ablate *Shh* only in the noto-chord and floor plate. **C.** An E18.5 *Foxa2-CRE-ERT2;Shh*^*-/fl*^ neonate after a Tamoxifen injection at E7.0 exhibits a similar phenotype to *Shh* null animals. Neonates develop without vertebrae and proximal ribs (inset – dorsal view). Distal ribs develop with mild segmental defects. **D.** The additional loss of a *Casp3* in the *Shh* hypomorph worsened the phenotype. Note a reduction of the distal segment compared to the *Shh* hypomorph alone (inset – dorsal view shows lack of vertebrae and proximal ribs). **E.** Normal embryo, ventral (top) and dorsal view (bottom). **F-H.** Range of hypomorph phenotypes within a litter. Those with the least severe phenotype **(F)** have some disruption of the vertebrae and the most proximal ends of the proximal ribs, while the most severe phenotype is similar to *Shh* KO embryos with complete loss of vertebrae and proximal ribs (**H**). Phenotypes were consistently observed in at least three animals of each genotype.

### Somite patterning does not differ between the *Shh* KO and *Shh;Apaf1* DKO embryos

It is well-established that somite patterning is specified by instructive signals from the surrounding tissues (Brent and Tabin, 2002). Specifically, *Shh* from the ventral midline is important for the induction and maintenance of sclerotome markers (Fan and Tessier-Lavigne, 1994; Fan et al., 1995; Furumoto et al., 1999; Marcelle et al., 1999). To determine if altered somite patterning could account for the more severe skeletal phenotype observed in the *Shh;Apaf1* DKO animals, we assessed the expression of markers for different somite compartments in control, *Apaf1* KO, *Shh* KO, and *Shh;Apaf1* DKO embryos. Compared to control embryos, *Apaf1* KO embryos showed no disruptions in overall somite, sclerotome, or dermomyotome patterning as assessed by RNA *in situ* marker expression analysis (data not shown) When examining *Shh*;*Apaf1* DKO embryos, we first found that they were smaller (compare panels 2G’, H’) and consistently delayed at E9-E12, (∼3 fewer somites than average for that litter; n > 6 litters) compared to *Shh* KO embryos suggesting the *Apaf1* plays a role in embryo size in the absence of *Shh*. Thus to compare patterning without size as a variable, embryos were carefully stage-matched by somite count (sometimes from different litters).

First we ruled out that the difference between *Shh* KO and *Shh;Apaf1* DKO embryos is due to defects in myotome development which could secondarily impact the sclerotome. Indeed, embryos null for skeletal muscle genes also display skeletal abnormalities, including rib fusions, truncations and sternal defects (*Pax3*: ((Henderson et al., 1999; Vivian et al., 2000)). In *Shh* KO embryos epaxial progenitors are reportedly absent, while the hypaxial progenitors are reduced (Borycki et al., 1999; Gustafsson et al., 2002) and our results looking at *Myf5* and *MyoD* expression confirmed this observation (Fig. 4A2, B2). Perhaps loss of the epaxial compartment explains loss of the proximal rib in both situations while additional defects in the hypaxial compartment are responsible for lost distal segments in the *Shh;Apaf1* DKO embryos. However, we found that *Shh;Apaf1* DKO embryos displayed the same pattern as *Shh* KO embryos (Fig. 4A3, B3) suggesting that a failure to specify the hypaxial muscle compartment is not the cause of the *Shh;Apaf1* DKO phenotype.

**Figure 4:**
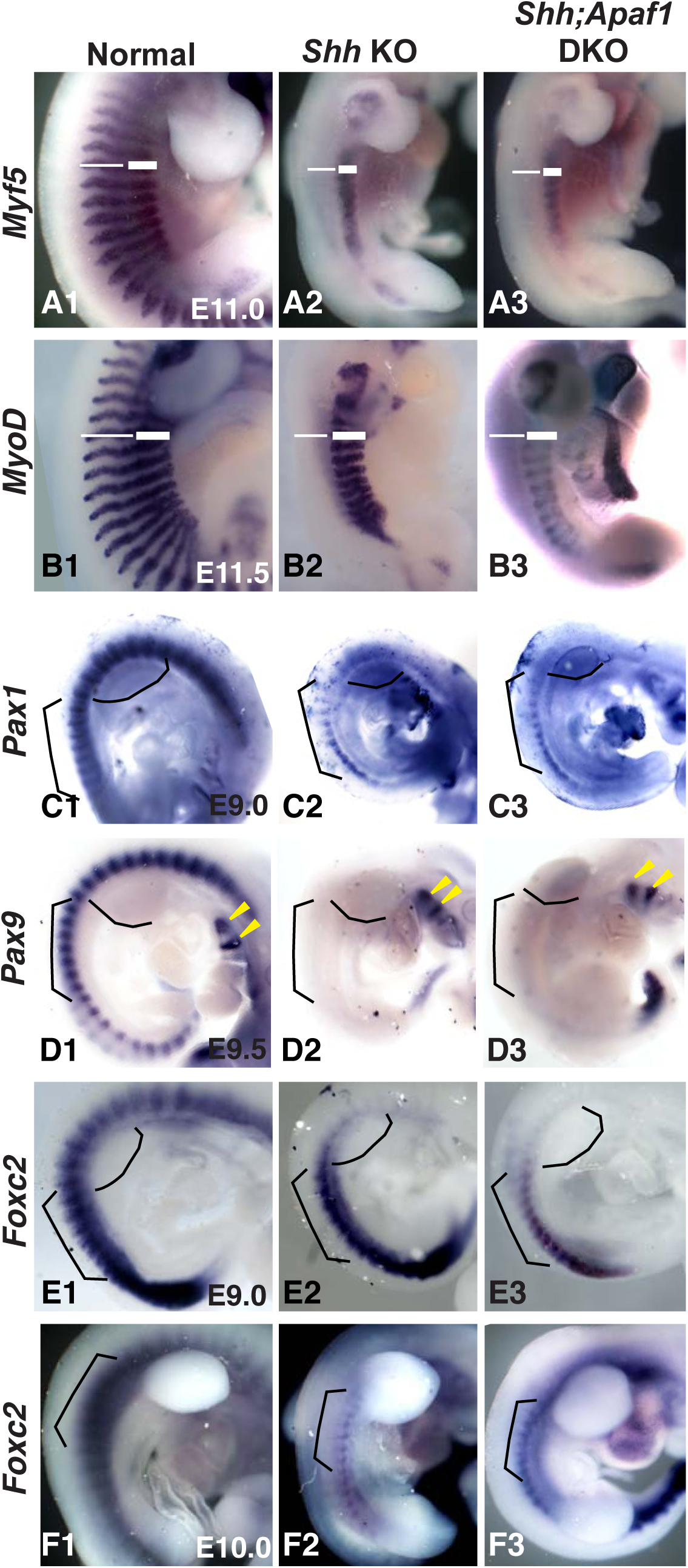
Somite patterning does not appear grossly altered when *Apaf1* is removed. **A1-B3**. E11.0 and E11.5 embryos showing *Myf5* (**A1-A3**) and *MyoD* (**B1-B3**) expression in the developing epaxial and hypaxial muscles. **A1 and B1.** Normal embryos. Loss of Shh leads to a lack of *Myf5* (**A2**) and *MyoD* (**B2**) in the epaxial region (thin line) but not the hypaxial region (thick line). *Shh;Apaf1* DKO embryos also display a loss of *Myf5* (**A3**) and *MyoD* (B3) in epaxial but not hypaxial progenitors. **C1-C3.** Analysis of *Pax1* expression at E9.0. **C1.** *Pax1* expression in normal embryos. **C2.** *Pax1* expression in the sclerotome of *Shh* KO embryos. **C3.** *Shh;Apaf1* DKO embryos also express *Pax1* in a similar pattern. **D1-D3.** Analysis of *Pax9* expression at E9.5. **D1.** *Pax9* is expressed in the sclerotome of normal embryos. **D2**. *Shh* KO embryos do not have *Pax9* expression in the sclerotome. Note *Pax9* expression in the brachial arches (arrowheads). **D3.** *Shh;Apaf1* DKO embryos exhibit a similar pattern. **E1-F3.** Analysis of *Foxc2* expression at E9.0 and E10.0. *Foxc2* is expressed throughout the developing somite at E9.0 (**E1**) and is restricted to the sclerotome by E10.5 (**F1**) in normal embryos. *Foxc2* expression is detected in similar patterns in both the *Shh* KO (**E2, F2**) and the *Shh;Apaf1* DKO (**E3, F3**) embryos at both E9.0 and after de-epithelialization of the thoracic somites (E10.0). Brackets outline comparable somite regions, curved line indicates forelimb bud location. Phenotypes were consistently observed in at least three animals of each genotype.

Another possibility is that in *Shh;Apaf1* DKO animals, sclerotome is never specified leading to the loss of both vertebral and sternal rib elements. *Pax9* and *Pax1* are expressed in similar patterns in the sclerotome and are required for rib and vertebrae development (Furumoto et al., 1999; Peters et al., 1999; Rodrigo et al., 2003). Previous studies have shown that *Pax1* is initially expressed in the *Shh* KO embryo but then lost (Chiang et al., 1996; Borycki et al., 1998; Zhang et al., 2001). As previously reported, we found *Pax1* expressed in a smaller domain in *Shh* KO embryos compared to normal or *Apaf1* KO embryos at E9.0 (Fig. 4C2). However, in *Shh;Apaf1* DKO embryos, *Pax1* expression was similar suggesting that some sclerotome specification still occurs (Fig. 4C3). As development continues, *Pax1* expression is eventually lost in *Shh* KO embryos as well as in the *Shh;Apaf1* DKO embryos (data not shown). Neither *Shh* KO nor *Shh;Apaf1* DKO embryos exhibited *Pax9* expression in somites at E9.0 and E9.5 (data not shown; Fig. 4D2-D3).

We then determined if using a more comprehensive sclerotome marker would produce the same results. *Foxc2,* along with *Foxc1,* are winged helix/forkhead transcription factors required for axial skeletal development with overlapping expression in the somites (Kume et al., 2001; Wilm et al., 2004). *Foxc2* is expressed early in the unsegmented paraxial mesoderm but then becomes restricted to the sclerotome, and may be a more comprehensive sclerotome marker than *Pax1/9* (Fig. 4E1, F1), (Furumoto et al., 1999). Like *Pax1/9*, prior to E9.0, *Foxc2* was expressed in a similarly reduced domain in both *Shh* KO and the *Shh*;*Apaf1* DKO embryos compared to normal embryos (Fig. 4E2, E3). However in contrast to *Pax1/9*, the smaller domain of *Foxc2* expression continued to be present after E10.0 in *Shh* KO embryos. We found that this was also the case in somite-matched *Shh;Apaf1* DKO embryos (Fig. 4E3, F3). In summary, in both contexts, the loss of *Shh* correlates with smaller *Pax1/9* and *Foxc2* expression domains. In addition, the *Pax1/9* domain disappears while the smaller *FoxC2* expression domain persists, suggesting that at least a portion of the sclerotome is maintained in both contexts. Thus, an alteration in early somite patterning does not readily explain the difference in the final skeletal phenotype between *Shh* KO and *Shh;Apaf1* DKO animals.

### Progenitor cells are specified but are unable to differentiate

We next determined if it is the failure of this remaining sclerotome compartment to undergo differentiation that distinguishes *Shh* KO from *Shh;Apaf1* DKO animals. Alcian blue staining detects the acid polysaccharide-rich matrix found in maturing cartilage. In control animals at E12.5, alcian blue stain in the presumptive rib cartilages extends dorso-laterally less than halfway around the chest (likely representing the proximal rib segments). In contrast *Shh* KO and *Shh*;*Apaf1* DKO embryos did not exhibit any signs of cartilage development in the thoracic region at this stage (data not shown). In control embryos a day later (E13.5), both proximal and distal cartilage anlagen now extended more ventrally towards the sternum (Fig. 5A). *Shh* KO embryos at this stage displayed some cartilage development, however the pieces were much smaller and located distally (aligned under the forearms) most likely representing the distal-most sternal ribs (Fig. 5B, B’). In contrast, no rib cartilages could be detected in *Shh*;*Apaf1* DKO embryos at E13.0 (Fig. 5C, C’). Thus, cartilage fails to mature in *Shh* KO animals when *Apaf1* is also removed.

**Figure 5:**
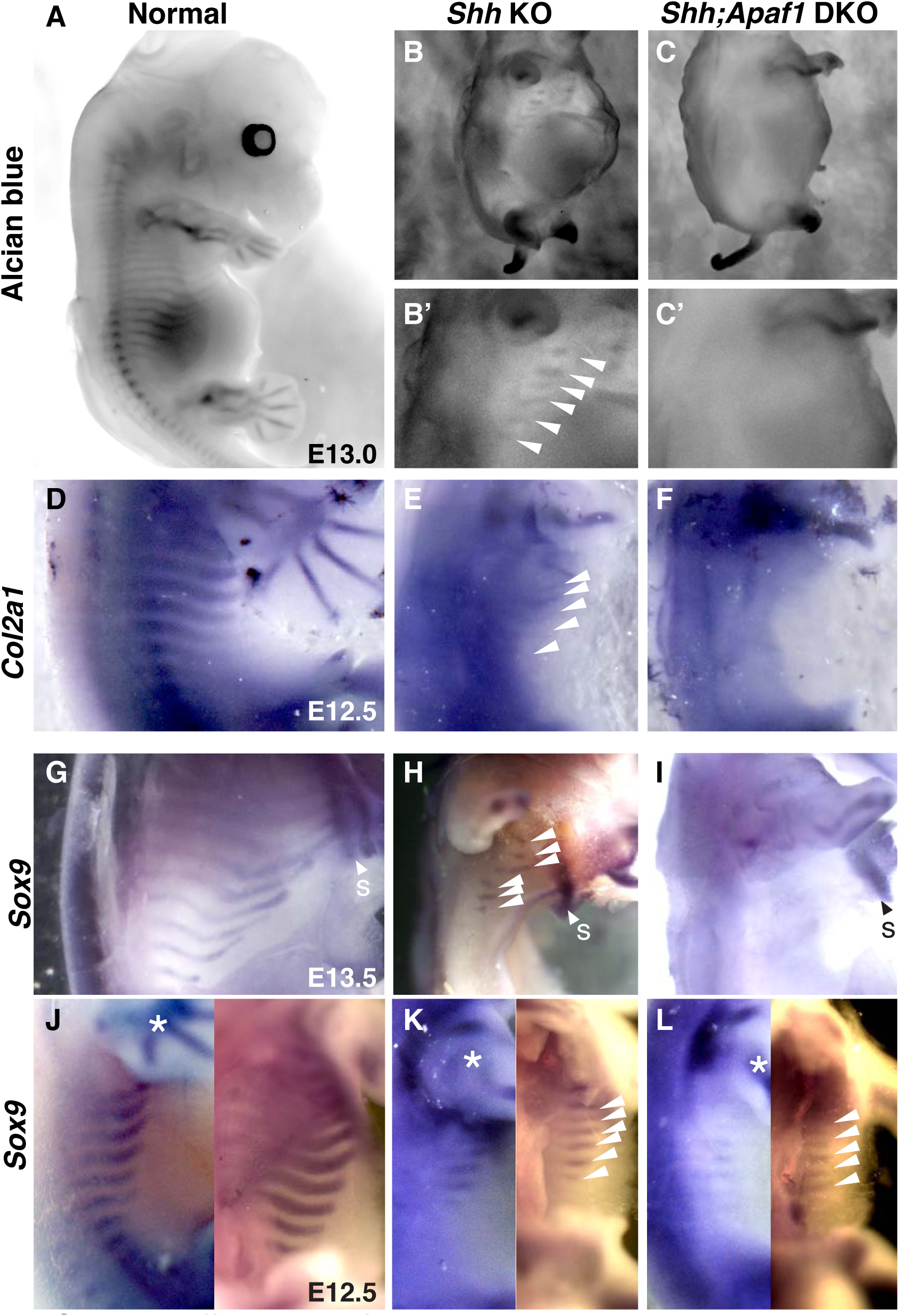
Cartilage differentiation fails in double null embryos. **A-C.** Alcian blue cartilage staining of E13.5 embryos is shown in black and white for greater contrast. **A.** Cartilage anlagen in normal embryos extend from the vertebrae towards the sternum. **B, B’.** *Shh* KO embryos have distally-located rib cartilages (white arrowheads). **C,C’.** No rib cartilages are observed in *Shh;Apaf1* DKO embryos. **D-F.** *Col2a1* expression at E12.5. **D.** Normal embryos express *Col2a1* in a band of cells from the vertebrae towards the sternum. **E.** *Shh* KO embryos express *Col2a1* only distally under the forearms. **F.** No *Col2a1* expression is observed in *Shh;Apaf1* DKO embryos where the ribs would develop. **G-L.** *Sox9* expression at E13.5 and E12.5. **G.** At E13.5, normal embryos express *Sox9* in the ribs extending from the vertebrae almost reaching the midline. **H.** In *Shh* KO embryos, *Sox9* expression is present but only distally. **I.** *Sox9* expression is undetectable in the entire thoracic area of *Shh;Apaf1* DKO embryos at E13.5. **J-L.** First panel shows an external lateral view, second panel an internal lateral view at E12.5, asterisk indicates *Sox9* expression in the limbs. **J.** Normal embryos express *Sox9* in the ribs extending from the dorsal midline toward the sternum. K. *Shh* KO embryos have reduced *Sox9* expression (thinner and shorter) and located only distally. **L.** *Shh;Apaf1* DKO embryos have further reduced *Sox9* expression in the distal domain. Phenotypes were consistently observed in at least three animals of each genotype.

Cartilage formation begins with the condensation of mesenchyme cells followed by chondrocyte differentiation. Perhaps the remaining sclerotome found in *Shh*;*Apaf1* DKO embryos is not able to differentiate into chondrocytes. Chondrocyte differentiation is observed through the formation of condensations that turn on *Collagen2* (*Col2a1*) and *Aggrecan* (*Agc*) (Lefebvre and Smits, 2005). At E12.5 in control embryos, *Col2a1* expression extends from the vertebrae laterally approximately halfway around the chest (Fig. 5D). *Shh* KO embryos exhibited *Col2a1* expression but only in a distal portion aligned under the forearms, most likely representing the chondrocytes for the distal-most sternal portion (Fig. 5E). However, no *Col2a1* expression was observed in the body wall of *Shh*;*Apaf1* DKO embryos (Fig. 5F). Examination of *Agc* at E13.0 showed a similar pattern (data not shown). Thus, loss of *Apaf1* along with *Shh* results in a failure in chondrocyte differentiation.

Sox9 is required for the specification of chondroprogenitors, chondrogenic condensation, differentiation, and proliferation (Akiyama et al., 2002). At E13.5, *Sox9* expression in control embryos is similar to the pattern of alcian blue staining and *Col2a1* expression. In *Shh* KO embryos, *Sox9* expression is found only in the distal compartment, while in *Shh*;*Apaf1* DKO embryos, *Sox9* expression is undetectable in the entire thoracic area, similar to the more differentiated markers, (Fig. 5H, I). We then looked earlier to determine if chondroprogenitors are *ever* evident. In control embryos at E12.5, *Sox9* expression is observed extending from the vertebrae laterally approximately halfway around the chest (Fig. 5J). At this stage, *Shh* KO embryos have *Sox9* expression but in a thinner reduced domain that is more distally located (likely representing the precursors of the distal-most sternal ribs). Interestingly, at E12.5, *Sox9* expressing cells are indeed present in *Shh*;*Apaf1* DKO embryos, however they are found in an even thinner and shorter domain. Quantification of this domain showed that *Sox9* expressing cells occupy approximately half the region compared to *Shh* KO embryos (Fig. 5K, L). This suggests that in *Shh*;*Apaf1* DKO embryos, chondroprogenitors are specified from whatever sclerotome is remaining, however in contrast to *Shh* KO embryos they are not maintained and fail to condense or differentiate any further.

### An agent-based simulation is sufficient to explain the phenotypes

Programmed cell death occurs normally and is thought to prune and sculpt tissues during development (Baehrecke, 2002). In the absence of the cell death effectors *Apaf1* or *Casp3,* embryos exhibit a dramatic decrease in this normal cell death. As a consequence, in the brain, decreased cell death results in increased progenitor cell number, massive brain overgrowth (exencephaly), and perinatal lethality (Cecconi et al., 1998; Yoshida et al., 1998). In contrast, while both *Apaf1* KO and *Casp3* KO animals have decreased somite cell death, they have normal thoracic skeletons. One possibility to resolve this discrepancy is that increased somite cell survival is not important for thoracic skeletal development—impacting some other tissue or dispensable in some other way. Another possibility is that in the context of the somite, compensation occurs and a sufficient number of progenitors for normal skeletal development is generated. For instance, perhaps in the absence of *Apaf1*, cell proliferation is decreased and thus the final number of cells needed to build the mouse thoracic skeleton is maintained. To determine if this kind of compensation might be occurring, or if some other property could explain the observed phenotypes, we decided to build an agent-based simulation using NetLogo (Wilensky, 1999) to model rib outgrowth and patterning up to ∼E12.5.

To build the simulation, we incorporated six important causal processes. These included: a Hh signal with varying concentration, variable cell death and proliferation rates, a progenitor pool that could vary in number, boundaries as would be created by surrounding tissues, and the potential effect of local cell-cell communication. We proposed that these processes predominated and therefore composed the minimal set of factors to be considered. We then represented cells as NetLogo agents (“turtles”) and created an initial field of these agents randomly placed in a square to represent the location of progenitor cells within the somite. Based on our results with the inducible *Foxa2-CreERT2* experiments, we assumed that somite cells are responsive to Hh signaling in an early time window. The field of cells was then programmed to change through time within a defined rectangular space according to simple parameters: for example, the levels, diffusion, and concentration of a Hh signal could be controlled. In addition cell death rate and timing could be modulated along with cell proliferation and the number of agents in the initial field. A change in cell fate was simulated by changing the color of each “cell” to red (for proximal) or blue (for distal) at each time step with the decision determined by the amount of local Hh signal available and with an adjustable probability of conversion. To account for the reinforcement of fate by local cell-cell communication (the “community effect” (Gurdon, 1988)), each cell was programmed to evaluate the local distribution of red or blue cells at each time step, and to convert to the local majority color when a local super-majority of other-colored cells surrounded it. To simulate the shape of the outgrowing rib within the body wall, the cells were programmed to move outward as they proliferated based on their degree of crowding at each time tick, with cells only moving when their local crowding was sufficiently high and then also confined in the rectangular space. When the cells hit the far right-side end of the defined space, the clock was stopped to indicate the end of outgrowth (see Methods for more details). We adjusted the parameters to determine which combinations would simulate the phenotypic outcomes observed in our single and combination mutants. Our expectation was that the settings required would provide predictive insight into the mechanisms responsible for the actual phenotypes.

While the agent-based simulation represents a highly simplistic scenario compared to real cells that have a particular developmental history, a complex relationship with their environment, and specific migratory properties, we were impressed to find that this small set of six processes was both necessary and sufficient to produce the whole range of observable phenotypes. First, the simulation could represent the response to a Hh concentration gradient reliably with the percent of proximal red and distal blue cells dependent on the Hh dose and gradient steepness regardless of the initial number of cells within the defined field (Fig. 6A, B, Supplemental Fig. 1B). Interestingly, the generation of a highly distinct border between the two elements required cells to be programmed to sense the fate of local cells within their environment (the “community effect”) as predicted by early studies on muscle differentiation (Gurdon, 1988). Second, alterations in the rate of cell death or of proliferation, while profoundly impacting overall size, did not have a large effect on the ratios of cells that contributed to a proximal vs. distal element. The exception to this was if we extended the time during which very high rates of cell death occurs to cover most of the period of growth, this could result in high contributions to the proximal or distal element stochastically depending on initial conditions. This is reminiscent of situations where development occurs in highly adverse conditions and widely variable outcomes are observed. Finally, we found that because the simulation incorporated boundaries on growth, it could mimic aspects of size control. A more detailed discussion of this kind of agent-base modeling that includes factors beyond the scope of this study is in preparation. See Fig. 6C and Supplemental Figure 1A, C for simulation outcomes for all genotypes.

**Figure 6:**
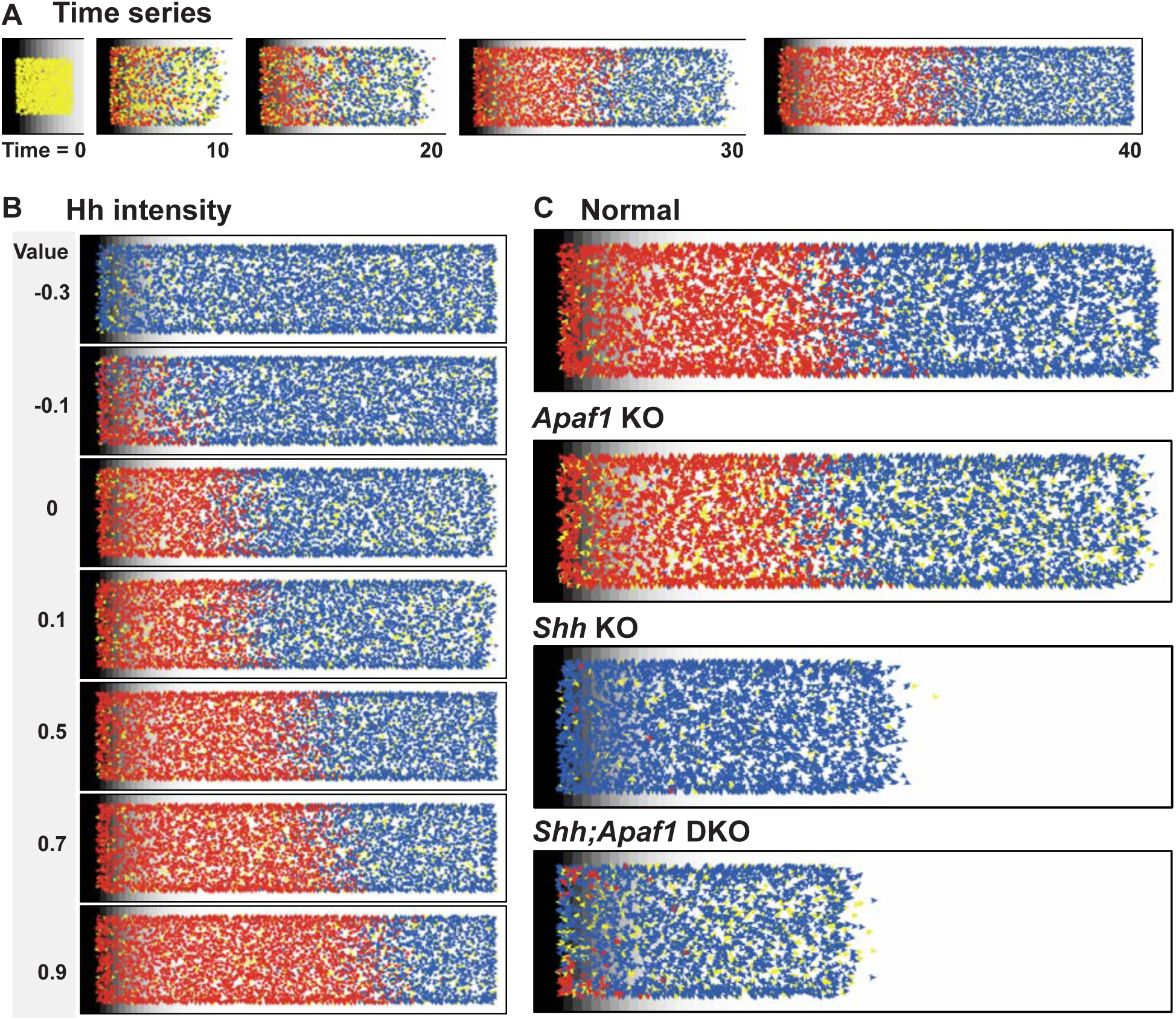
An agent-based model simulates rib proximal-distal patterning and outgrowth. **A.** A representative time series. The model simulates rib development up to E12.5 just prior to differentiation of the condensations. **B.** Outcome of adjusting Hh intensity without adjusting the gradient pitch. **C.** Representative outcomes that mimic the phenotypes. Normal was simulated with a moderate Hh gradient intensity, mild cell death, and a moderate proliferation rate. The loss of *Apaf1* was simulated by decreasing cell death to zero and moderately decreasing the number of initial progenitors; loss of *Shh* was simulated by decreasing the Hh intensity, decreased progenitor pool size, increased cell death. The absence of both Shh and Apaf1 was simulated with no cell death, an even smaller initial pool was used (due to decreased sclerotome induction and decreased proliferation) and an overall decrease in cell proliferation. Example parameters used to generate these outcomes along with mulitple simulation runs can be found in Supplemental File 1A, B, C. Example simulations can be found in Supplemental File 2 and the code to run the simulations can be found in Supplemental File 3.

To summarize the observed phenotypes, *Apaf1* KO animals are largely unchanged throughout development compared to wildtype. In the absence of *Shh*, the proximal elements are missing while a hypoplastic distal portion develops. In the *Shh;Apaf1* DKO somite patterning is similar to *Shh* KO embryos, but development is delayed. No proximal elements form, the distal elements are quite hypoplastic at E12.5 and then fail to differentiate.

To understand the role of Apaf1, we first used the agent-based simulation to compare the normal wildtype situation to the *Apaf1* KO situation. We found that if we mimicked the loss of *Apaf1* by decreasing the cell death rate and thereby eliminating normal cell death, we could easily replicate the normal *Apaf1* KO phenotype as long as the proliferation rate was also decreased. A decrease in the initial size of the progenitor pool could also contribute to replicating the phenotype.

Next we compared the *Shh* KO situation to wildtype. We found that decreasing the Hh concentration changed the proportion of blue vs. red cells contributing, with now mostly distal cells making up the final structure, similar to the observed phenotype of *Shh* KO embryos. A mild decrease in Hh levels could also result in an outcome similar to the hypomorphic phenotypes observed in the *Shh* KO conditional nulls (reduction in the proximal element). However, matching the observed decrease in size required including more cell death (as seen in Fig. 2), decreasing proliferation, or specifying that fewer initial cells are recruited to the rib anlage or some combination of these effects.

To distinguish between the different scenarios we mimicked the *Shh;Apaf1* DKO. Assuming an independence of the role of Shh and Apaf1, we set the Hh level to the same as used for the *Shh* KO. We also reduced the cell death to 0 and decreased proliferation rate to the level used in the *Apaf1* single KO. We found that this produced the correct distal pattern but a much larger rib than actually observed at E12.5. In order to match the observed decrease in size, either the initial number of cells must be decreased or the proliferation rate is decreased even further than as designated in the *Apaf1* KO simulation. These simulations indicated that in order to sufficiently explain the phenotypic outcomes, it was essential that we determine if the single and combination mutants exhibited changes in proliferation rate or in the number of cells initially participating in rib development, or both.

### *Apaf1* is important for determining somite size and proliferation rate

To analyze growth features and further constrain the range of possibilities suggested by the simulations, we collected somite-matched E9.0 embryos when multiple somites are readily visible and the forelimb bud has begun to emerge thus providing an internal landmark for analysis across all genotypes. Embryos were sectioned and select sections through the center of the somites were analyzed for size parameters and the expression of phosphorylated histone H3 (pHH3), an indicator of cells in mitosis. The number of cells were counted in the somite area, and nuclear density were measured to assess size (DAPI-positive area on cross-section) (Fig. 7A-D). We first found that while *Apaf1* KO embryos were indistinguishable from their normal littermate controls in terms of overall size, the somites were reduced in cross-sectional area by 25% (Fig. 7E). In addition, as predicted by our simulations, embryos lacking *Apaf1* did have a 48% lower proliferation rate compared to controls (Fig. 7F). Thus we suggest that a reduced cell death rate compensated by a reduced initial size and proliferation rate leads to normal outgrowth and patterning in the *Apaf1* KO.

**Figure 7:**
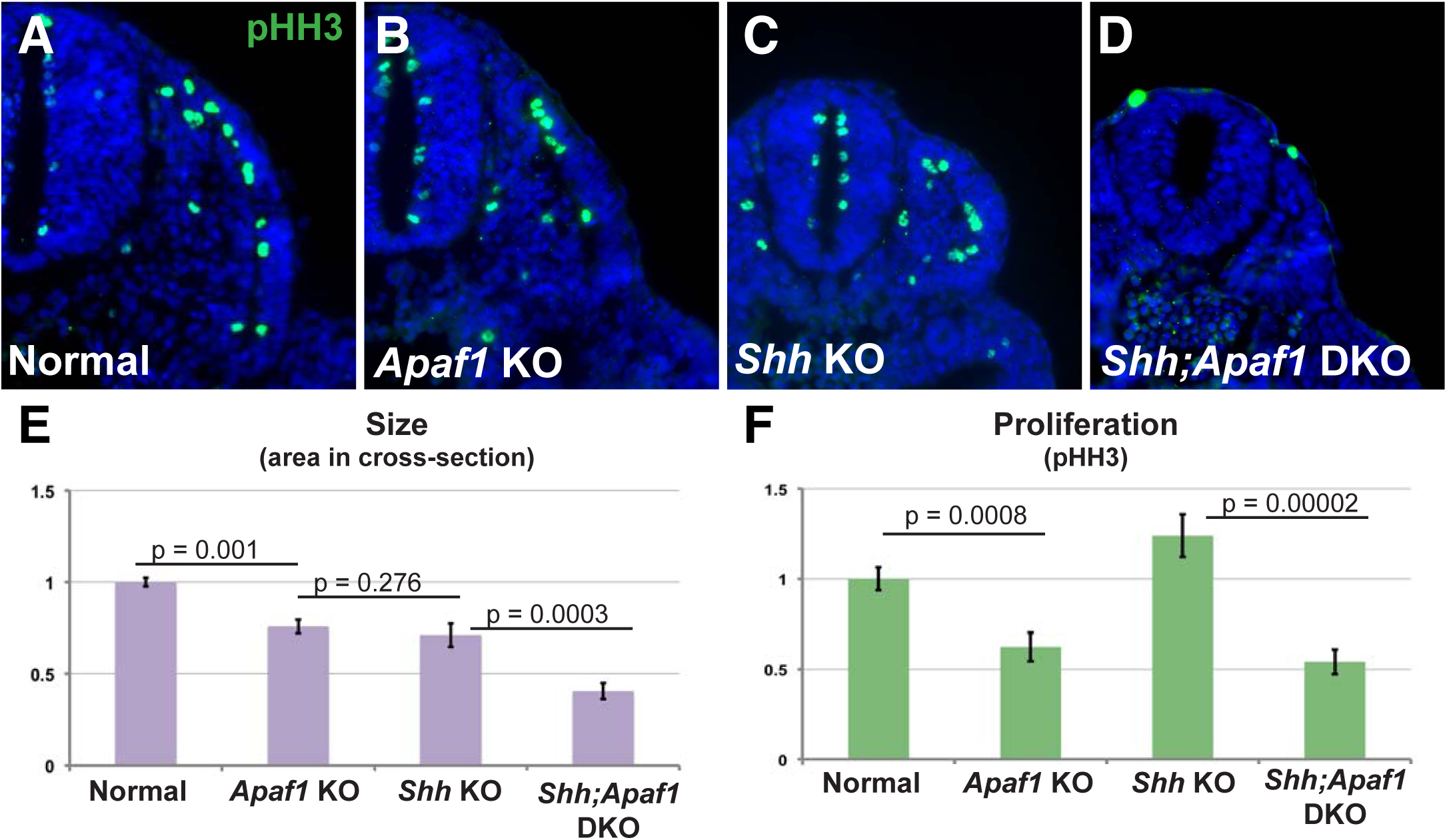
Alterations in somite size and proliferation are evident in the different genotypes. **A-D.** Representative transverse sections from somite-matched E9.0 mouse embryos at equivalent locations (based on anatomical landmarks). Phosphorylated histone H3 (pHH3) immunofluorescence (green) reveals cells in mitosis (blue: nuclei, DAPI). **E**. Somite size is measured by the cross-sectional area that is DAPI positive relative to normal. Shh;Apaf1 DKO embryos has much smaller somites (p = 2.52 × 10^−8^; other p values as indicated). **F.** Graph showing mitotic cells within the somite as a percentage of all nuclei in the somite. Normal and *Shh* KO embryos have a similar ratio of pHH3 positive to total cells in the somite (shown relative to controls, measured percentages = 6.1% +/- 0.4 vs. 7.4%+/- 0.7, difference not statistically significant, p= 0.110). While *Apaf1* KO (shown relative to controls, measured percentage: 3.8% +/- 0.5%) and *Shh;Apaf1* DKO (3.3% +/- 0.4) embryos both exhibit a decrease in the ratio of pHH3 positive to total cells. At least 2 animals per genotype were analyzed, 6-8 somites per animal; brackets = s.e.m). See also Figure 7—source data 1.

Loss of *Shh* resulted in a proliferation rate that was not greatly affected. Thus, the possibility that the smaller segment size is caused by reduced proliferation rate does not appear to explain the phenotypic outcome. The remaining possibilities generated by our simulations were some combination of increased cell death and decreased initial size of the progenitor population. Interestingly, both of these features could be observed in cross-section (Fig. 2 and 7E, about 30% smaller vs. normal controls).

Finally we considered the predicted possibilities for the *Shh;Apaf1* DKO and ruled out the possibility that a severely decreased proliferation rate (even further reduced from that in the *Apaf1* KO) accounts for the smaller somite size as the observed rate was essentially unchanged (Fig. 7F). The remaining possibility —an early reduction in the initial size of the progenitor pool due to the combination of a decreased proliferation rate and a decrease in sclerotome induction (Fig. 7E)— appears to be the cause of the further size reduction of *Shh;Apaf1* DKO vs. *Shh* KO rib condensations. Thus, despite reduced cell death, patterning is similar to the *Shh* KO as evident by RNA *in situ* hybridization but reduced somite size, delayed development and ultimately reduced condensations at E12.5 leads to failed skeletal differentiation at E13.5. Full animations and outcomes for all the phenotypes based on parameters inspired by these new data are available here: Supplemental File 1A,C, Supplemental File 2, and https://goo.gl/QwUE5B.

## DISCUSSION

In contrast to the vertebrate limb where multiple segments consisting of a number of skeletal elements form, proximal-distal patterning of the rib is comparatively simple with the formation of just two distinct skeletal elements. This simple situation combined with our observed alterations in proximal-distal patterning in different genetic contexts provides us with the opportunity to evaluate models of skeletal patterning in the context of growth. To understand how patterning arises, an important goal is to elucidate the mechanisms cells use to make decisions especially taking into consideration the information cells obtain from their local environment. Agent-based modeling, by virtue of placing the burden of computation on the cell (or “agent”), facilitates this thought process. Once a set of behaviors for individual cell decision-making and interaction is established, system-level patterns can then be observed to emerge from these interactions. This approach has been invaluable for understanding a number of important cellular phenomena such as quorum sensing among bacteria in biofilms (reviewed in (Gorochowski, 2016)), the arrangement of pigment cells into specific stripes in zebrafish skin (Volkening and Sandstede, 2015), and even how stem cell self-renewal could increase in developing mammary gland tissue that has been exposed to radiation (Tang et al., 2014).

In this study, we incorporate a role for Hh concentration in determining cell fate, the influence of cell proliferation and cell death on organ size, and the interaction of cells within boundaries in order to understand how a structure emerges from local decisions. By simulating a biological system using rules that respect local decision-making, and restricting the parameters to those compatible with the observed phenotypes in our genetically modified embryos, we infer a minimum set of rules that can be used to explain how pattern arises. We show that patterning can happen early when cell numbers are small and thus the distances required to interpret a Hh gradient are not too great. In addition we infer how size is regulated via mechanisms that control cell number such as initial progenitor number and proliferation rate. Through this analysis we have generated an overall picture for how patterning occurs that can be used to explain the different phenotypes observed (Fig. 8).

**Figure 8:**
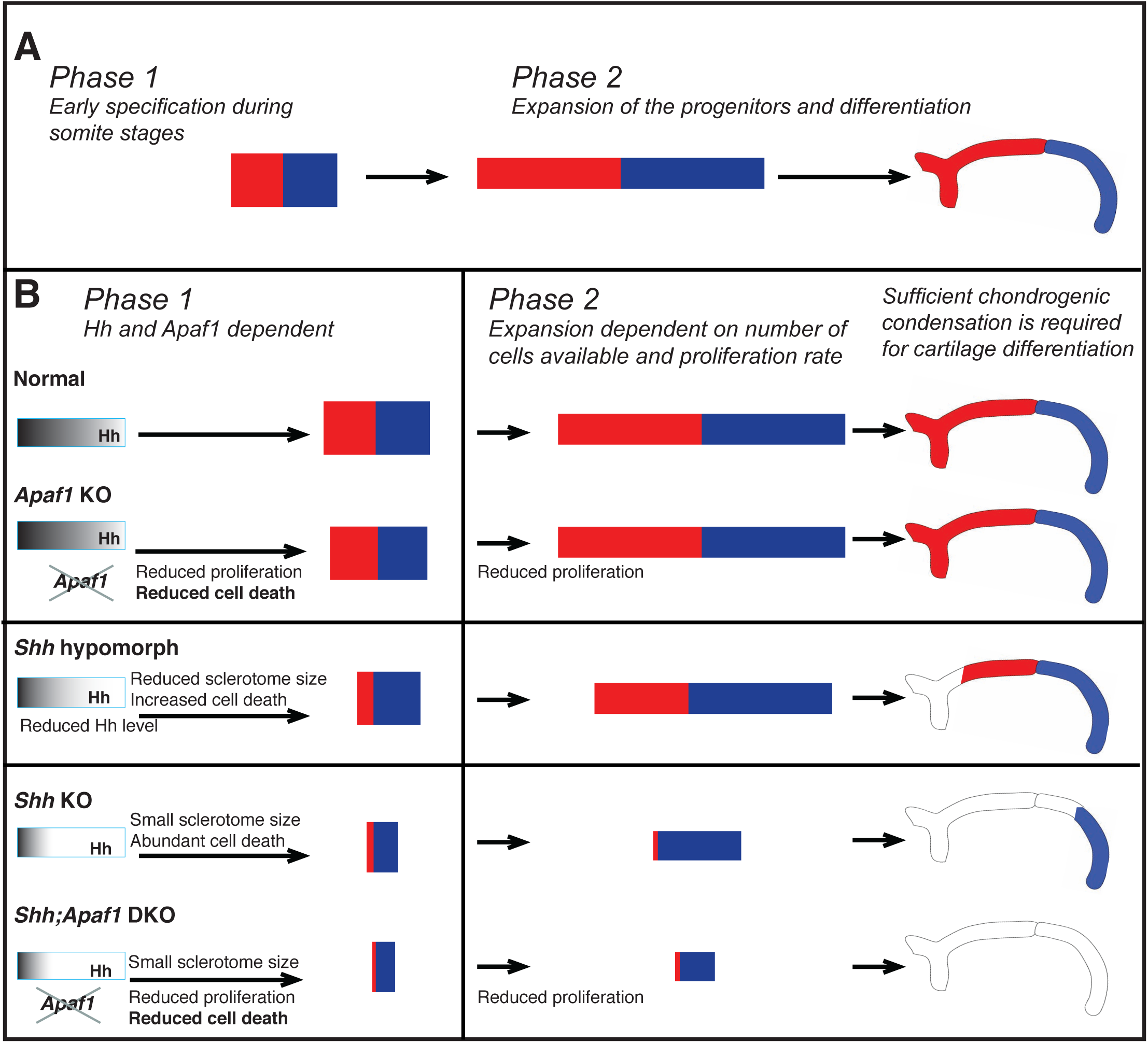
Two-phase model for rib proximal-distal patterning: early specification and expansion. **A.** We found that an early specification and expansion model could most readily explain the different phenotypes given our marker analysis, cell death, and proliferation data. This model proposes that the proximal and distal rib segments are specified early (during somitogenesis), (Phase 1) and then differentiation and expansion occurs during embryo growth (Phase 2). **B.** We suggest that during normal development, Phase 1 involves sclerotome induction, cell death, and proximal-distal patterning (all involvling a role for Hh), while Phase 2 involves expansion of the number of cells available to form both rib segments dependent on: 1) the events of Phase 1, 2) the proliferation rate, and 3) the constraints of the space in which expansion happens (body size). We propose that in the absence of *Apaf1,* progenitor specification happens normally as Hh signaling is not altered. The initial progenitor pool is slightly smaller due a decrease in cell proliferation but this is mostly compensated for by decreased cell death and normal development proceeds. In *Shh* KO and *Shh:Apaf1* DKO embryos, Hh signaling is greatly reduced during Phase 1 and thus the number of cells induced to become sclerotome is reduced (much smaller initial size). In addition, low Hh signaling may only be sufficient to specify distal sclerotome while high Hh is needed for complete proximal specification. In the *Shh* KO, despite a high level of cell death, evidently a sufficient number of cells proliferate and are able to differentiate into the most distal segment during Phase 2. When Hh signaling is not as dramatically reduced (as in *Shh* hypomorphs) distal and some proximal specification can occur. In *Shh:Apaf1* DKO embryos, however, along with reduced sclerotome due to both loss of *Shh* (smaller initial size), cell proliferation in the somite is also significantly reduced. Thus, despite the lack of cell death, there are ultimately overall fewer skeletal progenitors generated during Phase 2. These embryos are still able to form small distal condensations however they are smaller than those found in *Shh* KO embryos and thus may not contain a sufficient number of cells for cartilage differentiation to proceed.

### The role of Hh signaling is multi-functional and early

As observed in many other contexts (Briscoe and Small, 2015), the role of Hh signaling is multi-functional and likely required for many processes during rib development including: sclerotome induction, maintenance of cell survival, and proximal-distal specification. The relative importance of these different roles may depend on context, especially considering what other pathways may be either active or compromised.

As shown in previous reports, Hh signaling is required for sclerotome induction since Shh is necessary and sufficient to specify the sclerotome and also necessary to maintain it (Fan and Tessier-Lavigne, 1994; Borycki et al., 1998; Marcelle et al., 1999). Furthermore, in the absence of the receptor *Smo,* sclerotome fails to form as indicated by lack of *Pax1* expression (Zhang et al., 2001). Based on the *Smo* null phenotype one might expect that no ribs would develop at all. However, in *Shh* KO embryos costal cartilages form and sclerotome markers are still expressed. The presence of these structures may be explained by the expression of *Ihh* which is found ectopically expressed in the foregut of *Shh* KO embryos (Zhang et al., 2001; Fogel et al., 2008) and this Hh source could explain the presence of some *Pax1* expression in *Shh* null embryos and its later loss (Fig. 4). Ihh signaling may be sufficient to induce *Pax1* but not enough to induce a normal size sclerotome domain; later the embryo gets larger and not enough signal may arrive to maintain it.

Hh signaling is evidently important for cell survival, since in the absence of *Shh*, the number of cells undergoing cell death in the somite is vastly increased. Therefore both cell survival and the number of cells induced to participate, together determine the size of the initial cell progenitor pool. The role of Hh signaling promoting cell survival is likely transient and early (prior to E9.0) as indices of cell death (LysoTracker and TUNEL) are highest at E9.0 and by E11.5 have decreased with staining only apparent in the most caudal less-differentiated somites. Thus because of the transient cell death, as long as the proliferation rate is normal or at least sufficient, the cell death rate may not have a significant impact on patterning (as seen in the *Apaf1* null embryos).

While we can’t unequivocally rule out other models for proximal-distal patterning, a role for high and low levels of Hh signaling specifying proximal and distal fates is supported by the skeletal pattern observed when Hh signaling is mildly compromised. In particular, the range of phenotypes observed in the *Foxa2*-CRE;*Shh* hypomorphs suggests that mild decreases in Hh signal results in partial reduction in the number of cells specified as proximal, while severe decreases in Hh signal results in massive reduction in the number of cells specified as proximal. In addition to the phenotypic evidence, our gene expression data is also supportive of this dose dependence as there appear to be sclerotome populations that are more and less sensitive to a Hh signal. Considering the smaller domain of *Pax1* and failure to maintain it in *Shh* KO animals along with the loss of the proximal rib, we suggest that the *Pax1*-expressing population makes a significant contribution to the proximal rib segment. *Foxc2* is likewise expressed in a smaller domain in the absence of *Shh*, however, *Foxc2* expression is not lost later. We suggest that this remaining *Foxc2*-expressing population can be induced/maintained by low Hh signaling and represents cells that contribute to the distal segment which still forms in *Shh* KO embryos. How high vs. low Hh signaling specifies proximal vs. distal fates respectively is not clear but may involve different combinations of Gli transcription factors as has been elegantly demonstrated in neural tube patterning (Peterson et al., 2012; Cohen et al., 2015). Loss of *Gli2* and *Gli3* can influence the ability of the somite to respond to Hh signaling and turn on *Pax1* (Buttitta et al., 2003), however more studies will be needed to understand the details of this at the transcriptional level.

It remains an open question as to when proximal-distal specification occurs during early skeletal development. The two major competing hypotheses, based on studies in limb development, include the Early Specification Model and the Progress Zone Model (reviewed in (Mariani and Martin, 2003; Benazet and Zeller, 2009)). In the context of the rib, we favor an early specification scenario where proximal-distal specification occurs when the cells have not yet migrated long distances away from the midline and the progenitor pools are quite small. Therefore the transport of the Hh protein can occur over a short period of time and transport over long distances is not required to explain the results (Phase 1). Then as the embryo grows, the specified compartments would expand laterally (Phase 2; Fig. 7A). In recent years, this idea of early proximal-distal specification followed by expansion to establish the initial broad proximal and distal fields has been gaining traction in the field of limb development ((Rosello-Diez et al., 2011; Cunningham and Duester, 2015) and reviewed in (Mariani, 2010)) and seems easily applicable in the case of rib patterning. Support for a role for Shh signaling at very early somite stages (prior to E9.0), comes from our experiments where tamoxifen administration at E7.0 resulted in thoracic skeletal defects similar to *Shh* KO embryos (Fig. 3) while, administration after E8.0 or later was not sufficient to generate a thoracic phenotype. These results also align with previous studies suggesting that Hh signaling is only initially required at presomitic mesoderm stages and that subsequently BMPs are important in axial skeletal growth to maintain a chondrogenic regulatory loop (Zeng et al., 2002; Stafford et al., 2011).

In summary, there are probably several roles of Hh— influencing the number of cells induced to become sclerotome, reducing the number of cells undergoing cell death, and causing rib progenitors adopt a proximal vs. distal fate. The effect of Hh is likely early, during somite stages, and relatively short-range. Indeed within our agent-based simulations, the length scale over which Hh concentration can influence cell fate is approximately the same as the size of the progenitor pool and most cells “decide” their fate relatively early. We suggest that cells making late decisions become distal because they experience a very low Hh signal or become proximal due to influence from surrounding proximal cells.

### Apaf1 influences cell death and proliferation during rib development

Considering the prevalence of programmed cell death during development, including within the inner cell mass (Hardy et al., 1989), the relatively minor developmental defects found in the absence of apoptosis genes has been surprising. One explanation has been that genes in the apoptosis pathway can compensate for each other (Nagasaka et al., 2010) and/or that multiple genes need to be removed in order to see profound phenotypes. Another possibility is that cell death still occurs but by other cell death pathways (Yuan and Kroemer, 2010). However, another explanation is that the loss of these genes *does* block normal cell death and that the proper cell number is re-established by other mechanisms that decrease cell numbers such as decreased proliferation (Cecconi et al., 2004; Huh et al., 2004). In the context of rib development, with the absence of *Apaf1*, cell death is inhibited. When modeling this in our agent-based simulation, the proliferation rate must be moderately reduced to compensate for reduced cell death in order to achieve normal skeletal size. This prompted us to look more carefully at the proliferation rate and we did in fact see that it is decreased when *Apaf1* is lost (Fig. 7). We suggest that further perturbations (loss of *Shh*) influences the initial sclerotome cell number and this leads to the more severe defects seen in the *Shh*;*Apaf1* DKO embryos.

An interesting question for future research is determining the mechanism by which loss of *Apaf1* results in reduced proliferation. Two potential mechanisms are 1) the direct abrogation of the cell cycle machinery when the apoptosis pathway is blocked or 2) the indirect affect of a reduced number of dying cells that can release growth factors. Compensatory proliferation with dying cells releasing growth factors into their environment has been observed in other contexts (Fan and Bergmann, 2008b; Jäger and Fearnhead, 2012). Studies from *Drosophila* reveal that the signaling pathways involved in compensatory proliferation differ depending on the tissue and the developmental state of the tissue during stress. Highly proliferating tissues have been shown to induce Tgfβ and Wnt homologues (Pérez-Garijo et al., 2004; Ryoo et al., 2004), while differentiating tissues activate Hh signaling (Fan and Bergmann, 2008a). In the sclerotome, dying cells could be releasing growth factors that are sufficient to maintain the proliferation of the somite. In our studies, an increase in cell death in the *Shh* KO does correlate with a possible small increase in proliferation above normal (Fig. 7, not statistically significant). Then, in the absence of cell death (*Apaf1* KO), perhaps these growth factors are not released and this is a mechanism by which proliferation rate is decreased from normal.

### Skeletal development as biphasic

Based on our combined genetic analysis and agent-based simulations, it can be useful to think of rib development in terms of two main phases (Figure 8). During Phase 1 which starts as the presomitic mesoderm is forming into somites (E8.0-E10.0), the dominating activities are the induction of sclerotome and the establishment of proximal/distal cell identity mediated by Hh signaling while during Phase 2 (E10.0 and onwards) the dominating activity is expansion while cell identity is maintained. The transition between phases likely occurs gradually as the skeletal elements increase in size.

We propose that the ultimate size at birth of a skeletal element is determined by the number of cells left after Phase 1, along with an expansion of that population during Phase 2. Based on our agent-based simulations, the proliferation rate not only influences cell numbers early but, during Phase 2 can also have a profound impact on segment size because cell number increases exponentially. In the *Shh;Apaf1* DKO embryos, the decrease in transient cell death likely has a minor contribution compared to the decrease in sclerotome induction (due to loss of Hh signaling) and the decreased proliferation rate throughout somite development (due to loss of *Apaf1*).

### Final pattern is also influenced by the ability of progenitors to differentiate

Analysis of cartilage development in the different mutant contexts suggests that even when specification and expansion do occur, successful differentiation and maturation is needed to achieve the final skeletal pattern. In *Shh* KO embryos, a reduced distal domain forms that goes on to differentiate into small costal cartilages at E13.5. Whereas in the *Shh;Apaf1* DKO embryos, the reduced distal domain at E12.5 fails to maintain both *Sox9* and *Col2a1* expression at E13.5 indicating that differentiation has failed to occur (Fig. 5). While there could be some intracellular signaling mechanism by which *both* Hh signaling and Apaf1 are required to activate chondrocyte differentiation, another possibility is that chondrogenesis fails due to insufficient expansion. *In vitro* studies have shown that high cell-number and density is very important for chondrogenesis and for the production of cartilage matrix (Denker et al., 1999; Malko et al., 2013). Thus proper expansion of chondrogenic progenitors may be critical for differentiation and may be highly dependent on the density and numbers of progenitors. Decreased proliferation in *Shh;Apaf1* DKO embryos could result in a density of *Sox9*-expressing cells that is insufficient for differentiation to occur and thus a more severe phenotype develops. The requirement for a sufficient number of cells may also account for the variation in distal rib phenotypes seen in *Shh* KO embryos (Fig. 1).

Taken together, we suggest that during rib skeletal development, cells respond to local cues in a way that can be usefully modeled through a series of simple rules. During normal development, a compensatory feedback loop between the apoptotic pathway and cell proliferation plays a critical role in achieving the proper number of skeletal progenitor cells for cartilage differentiation. The precise molecular mechanism that underlies this balance may involve growth factor signaling but this is still to be discovered. Likewise, it will be important to determine if compensatory proliferation occurs in other embryonic contexts when programmed cell death is blocked. One of the most interesting issues for future investigation is to discover the molecular mechanism by which Hh concentration specifies proximal and distal identity and importantly the timing during which this happens. In addition, the precise mechanism by which refinement of pattern occurs is unknown and future studies could determine whether local signaling from the surrounding majority cells (community affect) plays a role or whether some other mechanism such as reciprocal inhibition or cell re-arrangement and assortment is involved.

Beyond the specifics of rib development, the use of mathematical and computational models can be extremely useful for making testable predictions. In particular, an agent-based approach allows the modeling of developing tissues while taking into consideration the behavior and fate decisions of individual cells based on local information. Furthermore, agent-based modeling is compatible with other classical differential equation-based modeling, such as reaction-diffusion and finite-element modeling which have been previously used to model limb development and other tissues (Zhang et al., 2013; Lau et al., 2015). In the future, these combined methods could be used to generate more precise multi-scale models (Zhu et al., 2010) which could be used not only to better understand the emergence of pattern in model organisms but also to understand how changes in cell behavior during evolution could generate new and diverse forms among different species.

## MATERIALS AND METHODS

### Generation of *Shh;Apaf1* and *Caspase3*;*Shh* DKO mice

To generate *Apaf1* and *Shh* double null embryos, females heterozygous for *Apaf1* and homozygous for a *Shh* ‘floxed’ conditional allele (*Apaf1*^-/+^; *Shh*^fl/fl^) (Yoshida et al., 1998; Lewis et al., 2001) were crossed with males ubiquitously expressing the CRE enzyme and heterozygous for both *Apaf1* and *Shh* (*β-actin-*CRE; *Apaf1*^-/+^; *Shh*^-/+^) leading to the production of *Shh:Apaf1* DKO embryos at a 1 in 8 frequency (12.5%). Heterozygous embryos were used as controls. A standard cross (*Shh*^*+/-*^*; Casp3*^*+/-*^ × *Shh*^*+/-*^*; Casp3*^*+/-*^) was established to create *Shh;Casp3* DKO embryos at a 1 in 16 frequency (6.25%)(Woo et al., 1998). To determine if Shh from the notochord and floorplate is required for rib development, conditional *Shh;Apaf1* DKO or *Shh;Casp3* DKO mice were produced using a Tamoxifen inducible *Foxa2-*CRE-ERT2 line (Park et al., 2008). Tamoxifen (Sigma; 3mg/mouse) and progesterone (Sigma; 1.5mg/mouse) were administered by intraperitoneal injection at E6.5-E10.0. Analysis was performed on carefully stage-matched sets of embryos using somite number and anatomical features. All animal procedures were carried out in accordance with approved Animal Care and Use Protocols at the University of Southern California.

### Histology and gene expression assays

#### Bone and cartilage staining

Embryos were collected E12.5-18.5, and fixed in 95% EtOH. Skeletal preparations were carried out following a standard protocol (Rigueur and Lyons, 2014).

#### Sectioning

Tissue was cryo-protected and embedded in Tissue-Tek OCT compound in a peel-a-way embedding mold for frozen sectioning.

#### RNA *in situ* hybridization

Embryos were harvested and immersion-fixed in 4% paraformaldehyde. Digoxigenin-conjugated antisense riboprobes were prepared (Roche, cat# 11175025910) and whole-mount or section RNA *in situ* hybridizations were color developed using NBT/BCIP (Roche, cat # 11681451001) according to previously established protocols.

### Agent-based modeling

An Agent Based Model (ABM) was developed using the NetLogo system version 6.0 (Wilensky, 1999). The model operates by initially creating a field of agents “cells” (“turtles”) and then evolving this field through time according to simple rules. A varying concentration of secreted Hh is modeled by using a Gaussian type curve with peak at × = -10 and a variable width and height. The agents are initially randomly placed in a square block between x=0 and x=10 and y=+- 5. They divide with base probability of 0.05 per time step and die with base probability 0.05 per time step, and the death rate is multiplied by a variable-relative rate and a time-varying curve using a Gaussian with a variable-duration width in time. At each time step there is a probability to convert from yellow (unspecified) to red (proximal) or blue (distal) and this is based on the local concentration of Hh. Once converted (biased in fate), “cells” ignore the Hh concentration (mimicking the window of competence to respond), but at each time point, the local concentration of red or blue cells is assessed and the cell is programmed to convert to the local majority color when a local super-majority of other-colored cells surround it.

As the cells proliferate, the cells spread with a velocity proportional to the local concentration of cells in its vicinity (gradient of the smoothed pressure) as calculated on the NetLogo "patches" that the "turtles" move around on. Cells only move when their local concentration is sufficiently high to overcome a resistance to crowding, which has the effect of creating a surface tension or membrane like effect around the cells that holds them together. In addition there is a diffusion of each cell in time. The net effect is that the cell field expands as if inside a balloon or boundary until they hit the edge of the spatial field whereupon they are reflected back a small random amount, thereby confining them into an elongated field. When the cell field hits the far end of the elongated field the clock is stopped, otherwise evolution continues for the specified number of time ticks. In some situations, the developmental "checkpoint" effect caused by hitting the boundary of the field models the slowing of the growth rate that naturally occurs as structures form in development. Without this boundary effect the number of cells would grow exponentially in time and without bound.

By varying the initial cell field size, the Hh concentration in space, the time duration, the coefficients that determine yellow-red and yellow-blue conversion probabilities, the cell death intensity, the proliferation rate, and the duration of the cell death in time, we can model a variety of conditions. Code to run the simulations can be found in Supplemental File 3 and https://goo.gl/QwUEFB and run after downloading the free NetLogo program at: https://ccl.northwestern.edu/netlogo/.

### Cell proliferation and death

#### Cell death

Lysosomal activity (which correlates with cell death) was assayed with the dye LysoTracker-RED (Invitrogen) as demonstrated previously (Fogel et al., 2012). The LysoTracker staining patterns were confirmed with TUNEL analysis (*In Situ* Cell Death Detection Kit, Roche).

#### Proliferation and somite size

Anti-phospho-Histone H3 antibodies (Upstate Bio) were used in 1:500 concentration, followed by an Alexa Fluor® 488 goat anti-rabbit IgG (H+L) antibody (A11008) at a 1:250 concentration. To calculate proliferation rates in the thoracic somites, the number of pHH3-positive cells were counted and expressed as a percentage of the total number of DAPI-positive nuclei within the defined perimeter of the somite (based on morphology). Statistical analysis was performed in Excel using a two-tailed Student’s test to generate p values.

## Acknowledgements

We gratefully thank Gail R. Martin and Andrew P. McMahon for advice during the initial stages of this project, Cheng-Ming Chuong and Leonardo Morsut for comments on manuscript drafts, and Audrey Izuhara and Christy Furukawa for technical assistance. This work was supported by the University of Southern California (to F.V.M. and I.K.M) and by a CIRM training grant (to J.L.F.).

## Author Contributions

F.V.M. and J.L.F conceived of the project, designed the experiments, and wrote the manuscript. J.L.F carried out experiments with assistance from I.K.M. D.L.L. and F.V. M. designed the agent-based model.

The authors declare that there is no conflict of interest regarding the publication of this article.

